# Ketone body b-Hydroxybutyrate does not extend lifespan, but upregulates fecundity in food-limited *Daphnia*, with a transgenerational effect

**DOI:** 10.1101/2025.01.18.633735

**Authors:** A. C. Pearson, S. Bhadra, L. Y. Yampolsky

## Abstract

Ketone bodies accumulate on ketogenic diets and are known to have numerous beneficial health and longevity effects. Here we investigate the effects of exposure to environmental b-hydroxybutyrate (BHB), a ketone body, on longevity and fecundity of a model organism *Daphnia magna,* a plankton crustacean, maintained at limited food availability. We report that exposure to continuous or intermittent lifetime exposure to 2.5 – 10 mM of BHB reduces *Daphnia* lifespan, while intermittent exposure administered for 20-day periods has little effect on post-exposure survival, regardless of the age at exposure onset. On the other hand, various BHB exposure regimes significantly increased fecundity, including fecundity up to 100 days past a 20-day exposure. We further demonstrate that even relatively brief early life maternal exposure to BHB can increase daughters’ fecundity, although this transgenerational effect is genotype-specific. We argue that this effect must have a signaling nature rather than simply manifestation of additional source of energy provided by BHB and discuss potential significance of genetic variation in transmission of such signals.

## Introduction

Ketogenic diet, originally developed to remediate epilepsy (Ref) has been proven effective in promoting weight loss and longevity (Balietti et al. 2010; Edwards et al 2014; Campisi et al. 2019; Han et al. 2020; Wang et al. 2021) through a spectrum of mechanisms centered on energy source and signaling functions of ketone bodies – acetone, beta-hydroxybutyrate (BHB, also often abbreviated as βOHB) and acetoacetate – that accumulate in tissues when the organism consumes (and catabolizes) substantially more fats than carbohydrates (Newman & Verdin 2014; 2017; Veech et al 2017; Møller 2020; Madhavan & Stubbs 2025). BHB and other ketone bodies serve as non-sugar, lipid-derived energy source, particularly in the nervous system, and perform a variety of regulatory functions. As an energy source, ketone bodies differ from glucose in its energy metabolism propensity to increase in the NAD+/NADH ratio because synthesis of acetyl-coA from ketone bodies consumes more NAD+ molecules than synthesis of acetyl-coA as the final product of glycolysis (Newman & Verdin 2017). Additionally, all NAD+ consumed during this process are consumed in the mitochondria, thus preserving cytosolic NAD+, as opposed to half of NAD+ consumption that occurs in cytosol during glycolysis. Thus, generating similar mole per mole amounts of ATP, oxidation of ketone bodies maintains a higher NAD+/NADH ratio and therefore redox balance in cells and tissues, than oxidation of glucose. Additionally, oxidizing BHB increases efficiency of proton gradient during membrane phosphorylation relative to mitochondria utilizing products of glycolysis (Veech 2004).

Besides being an alternative energy source, BHB can also play a variety of signaling roles (Newman and Verdin 2014; 2017; Wang et al. 2021), including, notably, activation of NAD-dependent sirtuins by increasing NAD+ availability and inhibition of class I histone deacetylases (and hence maintaining elevated gene expression). NAD-dependents sirtuins are mitochondrial protein deacetylases (Stein & Imai 2012) that function as nutrient-responsive regulators and are thought to extend lifespan in yeast, worms and flies, and to be one of the underpins of caloric restriction extension of lifespan (Ghosh 2008; Kwon & Ott 2008; Sugishita et al. 2024.). Histone hyperacetylation induces expression of several key regulatory proteins, including Foxo3a, the mammalian ortholog of the stress-responsive transcriptional factor DAF16 that regulates life span in C. elegans (Kenyon 2010; Shimazu et al. 2013; Newman and Verdin 2017). Another downstream process affected by histone hyperacetylation is autophagy (Yi et al. 2012). Besides histones, BHB-inhibited deacetylases regulate a variety of other proteins, including NF-κB, TP53, and several other key regulatory proteins (Glozak et al. 2005). Besides histone hyperacetylation, BHB can also directly attach to lysins in histones and other proteins (post-translational protein β-hydroxybutyration; Xie et al. 2016), with additional not fully understood downstream transcriptional effects (Newman and Verdin 2017). Additional port-translational modification pathways are affected by BHB accumulation due to increase in generation of acetyl-coA and consumption of succinyl-coA, both of which are substrates for protein modification. Finally, BHB binds to and regulates a variety of membrane receptors, channel and transporter proteins. This plethora of regulatory activities, including those that directly affect NAD+/NADH ratio, antioxidant pathways, Foxo3a expression, and autophagy has laid foundation for the hypothesis that BHB may be the key intermediate in the caloric restriction benefit for longevity (Newman and Verdin 2014) and that ketogenic diet or extraneous administration of BHB can mimic the caloric restriction effect (Veech et al. 2017; Newman et al. 2017). Elsewhere (Pearson and Yampolsky, submitted) we test this hypothesis in an emerging model for longevity and aging studies, plankton crustacean *Daphnia* reporting that BHB exposure reduces early life mortality in *ad libitum* fed *Daphnia* to caloric-restricted level and results in intermediate pattern of gene expression. Here we ask the question of possible effects of BHB administration on *Daphnia* maintained at a low food level amounting to moderate caloric restrictio, aiming to see if a further extension of lifespan could be observed. Instead, we observe a moderate increase in fecundity, that extends, at least temporarily, trans-generationally, and discuss possible energetic and regulatory mechanisms of this effect.

Effects of ketone bodies on fecundity are poorly understood. In mammals, reproductive effort is controlled by metabolic fuels availability (Wade & Schneider 1992) and tissue concentration of ketone bodies is one of the signals indicating high energy supply, including energy from catabolizing body fats (Matsuyama & Kimura 2014). As the result, there is a significant literature on the role of ketone bodies in regulating fertility in cattle (Missio et al. 2022). Furthermore, in mammals, in which a significant portion of metabolic costs of reproduction occurs during lactation, increases concentration of BHB can have the opposite effect on further reproduction. For example, in cattle BHB injections decreased follicle growth, although did not alter ovulation regulation (Missio et al. 2022). Ketogenic diet is even being discussed as a fertility improving measure in human females (Kulak & Polotsky 2013), although clinical evidence is weak and mechanisms unclear.

*Daphnia* is an excellent model organism for the studies of longevity, reproductive allocation, and transgenerational effects. It reproduces by cyclic parthenogenesis, which allows maintaining genetically uniform, and yet outbred female-only lineages (referred thereafter as clones), and thus eliminating genetic heterogeneity in longevity and maternal effect studies. Transparent body allows direct in vivo measurement of fecundity and lipid allocation. Vast literature exists on physiology, aging, and effects of caloric restriction on longevity in this model (see Beam et al 2024 for a review) and genomic resources are available.

Maternal and transgenerational effects in *Daphnia* are also well characterized (LaMontagne, & McCauley 2001; Padilla Suarez et al. 2023; Agrelius & Dudycha 2025) and in the last few years there has been an explosion of publication making *Daphnia* one of the best studied organisms with respect to transgenerational effects. Maternal exposure to a variety of naturally occurring or anthropogenic environmental factors such as heat (Garbutt et al. 2014; Walsh et al. 2014; Lyu et al. 2017), hypoxia (Andrewartha & Burggren 2012), salinity (Mikulski & Mazurczak 2023), photoperiod (Toyota et al. 2019), UV light (Sha et al. 2020), radiation (Sarapultseva & Dubrova 2016), food availability (LaMontagne, & McCauley 2001; Ben-Ami et al. 2010; Stjernman & Little 2011; Garbutt & Little 2014; Coakley et al. 2018; Agrelius et al. 2023) and quality (Frost et al. 2010), population density (Michel et al. 2016), presence of predator cues (Walsh et al. 2015, 2016; Sha et al. 2020), presence of toxic or non-toxic cyanobacteria (Jiang et al. 2013; Dao et al. 2018; Radersma et al. 2018; Walsh & Gillis 2021; Zhu et al. 2024; Shahmohamadloo et al. 2024), presence of pathogens (Ben-Ami et al. 2010; Paraskevopoulou et al. 2022; Sun et al. 2023), as well as treatment with hormones (LeBlanc et al 2013), drugs (Michalaki & Grintzalis 2023), toxicants (Harney et al. 2022; Im et al. 2023; Lee et al. 2023; Zhu et al. 2024), or nanoparticles (Martins & Guilhermino et al. 2018; Qi et al. 2022) have been shown to elicit phenotypic changes in unexposed parthenogenic offspring or accumulating response in multigenerational treatments. Likewise, several studies reported the effects of maternal age on offspring life history traits (Sakwińska 2003; Plaistow et al. 2015; Anderson et al. 2022),) and one of them emphasized that even genetically uniform, identically treated, same age mothers may produce offspring that systematically differ in their life history (Sakwińska 2003).

Many of these studies focused on plastic (hermetic) increase in specific stress tolerance in the offspring of mothers exposed to the same pathogen (Ben-Ami et al. 2010; Paraskevopoulou et al. 2022) or the same stress (Mikulski & Mazurczak 2023). Perhaps the most striking example of hermetic maternal effect is higher lipid and protein provisioning (Guisande & Gliwicz 1993) and, predictbly, higher starvation tolerance (Gliwicz & Guisande 1992) and lower offspring food acquisition rate (Garbutt & Little 2014) in offspring of mothers experiencing low food conditions. Likewise, poorly fed mothers produce larger offspring (Boersma 1995; 1997; McKee & Ebert 1996). In many others the maternal stress and offspring response were orthogonal, such as for example, maternal temperature affecting parasite resistance (Garbutt et al. 2014), or, inversely, pathogen exposure affecting heat tolerance (Sun et al. 2023). Likewise, several studies reported transgenerational effects of population density, or food quantity or quality affecting offspring pathogen resistance (Frost et al. 2010; Stjernman & Little 2011; Michel et al. 2016). Another example of seemingly orthogonal environmental factor and transgenerational phenotype is the effect of cyanobacteria on *Daphnia* eye size (Walsh & Gillis 2021). Furthermore, in several cases environmental stress resulted in life history changes that were not necessarily adaptive or were maladaptive (Shahmohamadloo et al. 2024).

Thus, it appears that almost every possible environmental factor can elicit almost any kind of transgenerational change in *Daphnia.* Yet, one life history train stands out and it is fecundity. Irrespective the nature of maternal offspring environments, in many cases it was fecundity that responded along with, or even stronger than any other, presumably adaptive, life history or physiological traits (Paraskevopoulou et al. 2022; Lee et al. 2023), sometimes with an unexpected increase (Zhu et al. 2024). It appears, therefore, that fecundity changes, including fecundity increase, is often the preferred response to maternal signals about a wide variety of environmental changes. This rule is not universal, though, as some studies showed no transgenerational changes in fecundity (Shahmohamadloo et al. 2024), at least no such effect lasting through 2 unexposed generations, although even in this study fecundity did increase in response to *Microcistis* exposure in granddaughters of previously exposed grandmothers in 2 out of 8 genotypes, suggesting genetic variation for transgenerational effects, i.e. genotype-by-maternal environment interactions. Maternal effects for life-history traits apparent in some, but not other clones of *Daphnia* have been also observed elsewhere (e.g. Stjernman & Little 2011; Jiang et al. 2013; Michel et al. 2016).

Although in some cases maternal effects could be explained by direct influences of maternal phenotypes, for example via neonates’ body size (Garbutt & Little 2017) or lipids provisioning (Anderson et al. 2020), in most cases there is an indirect transgenerational epigenetic signal in action, particularly in the cases when the effect of maternal environment persisted for more than one unexposed generation. There is some evidence that at least some of these transgenerational effects is based on transgenerational changes in DNA methylation (Vandegehuchte et al. 2010; Jeremias et al. 2018; Feiner et al. 2022; Harney et al. 2022; Agrelius et al. 2023) or on histone modification (Lai et al. 2016). In contrast, there is no support for microRNAs’ role in transgenerational effects (Hearn et al. 2018). Either way, transgenerational effects are useful in pinpointing signaling mechanisms responsible to response to environmental stimuli and this is rationale for the inclusion of transgenerational experiment into this study.

## Materials and Methods

We report here results of four separate life-table experiments that included BHB expore as one of the treatments. These experiments are summarized in Table 1 and correspond to experiments 13, 23, 11, and 12 of Beam et al. 2024, respectively, where details on clones’ provisioning and maintaining and on general experimental protocol are provided. The experiments varied in handling details and BHB protocol, as described in Table 1. Briefly, *Daphnia* clones were maintained through parthenogenetic reproduction in modified ADaM medium (Klüttgen et al. 1994) at 20 °C and 12:12 L:D photoperiod, fed daily with *Scenedesmus acutus* culture. To collect experimental animals lineages originated from several progenitor females from each clone were maintained for 2 generation at the standard conditions of 1 daphnid per 20 mL with food added daily to the concentration of 10^5^ cells/mL and with water changed and neonates removed every 4 days.

**Table 1.** Summary of 4 experiments.

| Experiment | Set-up / group size | BHB treatments | Traits measured | Other treatments | Clones used<sup>4</sup> |
| --- | --- | --- | --- | --- | --- |
| Experiment 1 | 10 individuals per 50 mL insert, 1 insert per 200 mL bottle | 0, 5, 10 mM, lifetime, continuous. | Longevity, age-specific fecundity<sup>1</sup> | 2x food level | GB-EL75-69<br>IL-M1-8 |
| Experiment 2 | 50 individuals per 1 L insert, 4 inserts per 4 L tank, individually after age 100 days | 0, 2.5, 5 mM, lifetime, 2 hours per day, every day. | Longevity, age-specific fecundity<sup>2</sup> | NMN exposure<sup>3</sup> | GB-EL75-69<br>IL-M1-8 |
| Experiment 3 | 5 individuals per 50 mL, 1 insert per 100 mL bottle | 5 mM for 20 days at various ages; 2 hours per day, every day. | Longevity, age-specific fecundity<sup>1</sup> | NMN exposure<sup>3</sup> | IL-M1-8 |
| Experiment 4 | Mothers: as in Experiment 3; daughters: 1 individual 20 mL vial | Maternal 2.5 mM, 20 days at ages 10 – 30 days; 2 hours per day, every day. | Size at birth and at first reproduction, lipid content, and fecundity<sup>1</sup> (measured in daughters) |  | GB-EL75-69<br>HU-K-6<br>IL-M1-8 |

<sup>1</sup>Fecundity measured every other day as the number of offspring produced per day in each replicate jar, normalized by the number of females in the jar, and smoothened by sliding window of 3 consecutive datapoints.
<sup>2</sup>Fecundity measured as mean clutch size of individual females, including skipped clutches recorded as clutch size 0.
<sup>3</sup>NMN: nicotinamide mononucleotide. For results of this treatment see Pearson et al., in preparation.
<sup>4</sup>Clone IDs from Basel University Daphnia stocks collection (Basel, Switzerland). Clones will be thereafter referred to by the first 2 letters of clone ID.

Experimental animals (females only) were collected from these lineages within 24 h of birth and placed into control or BHB treatment groups as described in Table 1. Medium volume was adjusted at every water change to maintain the density of 1 daphnid per 20 mL. Feeding conditions (10^5^ cells/mL *Scenedesmus* per day, 1 daphnid per 20 mL of medium) were the same in all experiments reported here and correspond to moderately restricted dietary regimen (Beam et al. 2024; See Pearson et al., in preparation, for the effects of BHB exposure on life history of daphnids fed *ad libitum*). In Experiments 2 – 4 the *Daphnia* were housed in plastic inserts equipped with 1 mm mesh on the bottom which allow minimal handling during water change, neonate removal and BHB exposure (Cho et al 2022). Handling and housing of *Daphnia* was the same throughout the lifespan in all experiments except Experiment 2, in which the 1 L inserts do not allow sufficient medium level after the cohort size drops below 10 individuals. Therefore at the age of 100 days daphnids were moved from inserts into individual vials containing 20 mL of the medium. Fecundity data were analyzed separately for pre– and post-100 day switch in this experiment.

BHB exposure was either lifetime (Experiments 1 and 2) or limited to certain age classes (Experiments 3 and 4). Specifically, in Experiment 3, six cohorts were set up with the onset of 20-day exposure period ranging from 15 to 120 days; in 4 youngest of these cohorts there was still substantial fertility past the exposure to allow fecundity measurements. In Experiment 3 the single maternal cohort was exposed for 20 days between ages of 10 and 30 days. In Experiment 1 daphnids were exposed to BHB continuously, with fresh solution added at each water change, while in the other three experiments the exposure consisted of moving the mesh-bottom inserts containing daphnids from the housing jars into exposure trays containing BHB solution for 2 hours per day, every day. Control inserts were transferred for the same amount of time into identical trays containing ADaM medium. No food was provided during the exposure and the BHB solutions were filtered after each exposure to remove neonates and feces and stored at 8 °C and reused for no longer than 2 weeks. Such treatment eliminates bacterial growth in the medium with BHB, which may affect longevity and fecundity of the experimental animals.

In Experiment 4 the exposure to BHB was conducted in the maternal generation, while life-history parameters were measured in the daughters of exposed animals to measure maternal/transgenerational effects. Mothers were exposed as described above for the first 30 days of their life and the offspring born during exposure period or 4 days past the exposure period were discarded, to ensure that the daughter generation individuals were exposed to BHB as germline cells / oocytes and not during embryonic development. These individuals were used to either measure size and lipid content at birth by photographed under a fluorescent microscope after 2 hours 1 ug/mL Nile Red exposure, see Anderson et al., 2022 for details).

In experiments in which daphnids were kept in groups of 5 or 10 in 100 or 200 mL jars, i.e., Experiments 1 and 3, fecundity was measured every other day as the number of offspring produced per day in each replicate jar, normalized by the number of females present in the jar at the start of the 2-day period, and smoothened by sliding average with the window of 4 days (e.g. over 2 consecutive datapoints). In Experiment 2, six females were randomly sampled, with replacement, from each of replicate tanks and eggs in their brood chambers were counted, including any skipped clutches recorded as clutch size 0. Finally, in the final stage of Experiment 2, and in the daughter generation of Experiment 4, in which females were maintained in individual vials, all offspring produced by each female were recorded. Thereafter, fecundity measured as a sliding window average of neonates produced per female will be referred to as sliding window fecundity, while fecundity measured as clutch sizes of individual females will be referred to as clutch size.

Survival and longevity data were analyzed by Proportional Hazards model with clones, BHB treatment and their interaction as factors. Fecundity data were square root-transformed for normality. Fecundity, body size at birth and maturity, and lipid fluorescence in neonates measured in the daughters’ generation in Experiment 4 were analyzed using REML analysis of variance, with the same factors as main effects and with maternal ID, nested within clones, as a random effect.

## Results

### BHB exposure did not extend longevity in food-limited Daphnia

Contrary to expectations, life-long exposure to 2.5 – 10 mM BHB reduced lifespan in either of two clones tested (Experiments 1 and 2, Fig. 1). The reduction effect did not appear to depend on whether the exposure was continuous or intermittent (Fig. 1 A,B vs. C.D) and did not show dosage effect in either experiment (5 vs. 10 mM or 2.5 vs 5 mM). In Experiment 2 the longest-surviving BHB-exposed individual outlived their control counterparts, but this maximal lifespan extending effect did not counterbalance generally higher mortality earlier in life.

**Fig. 1.**
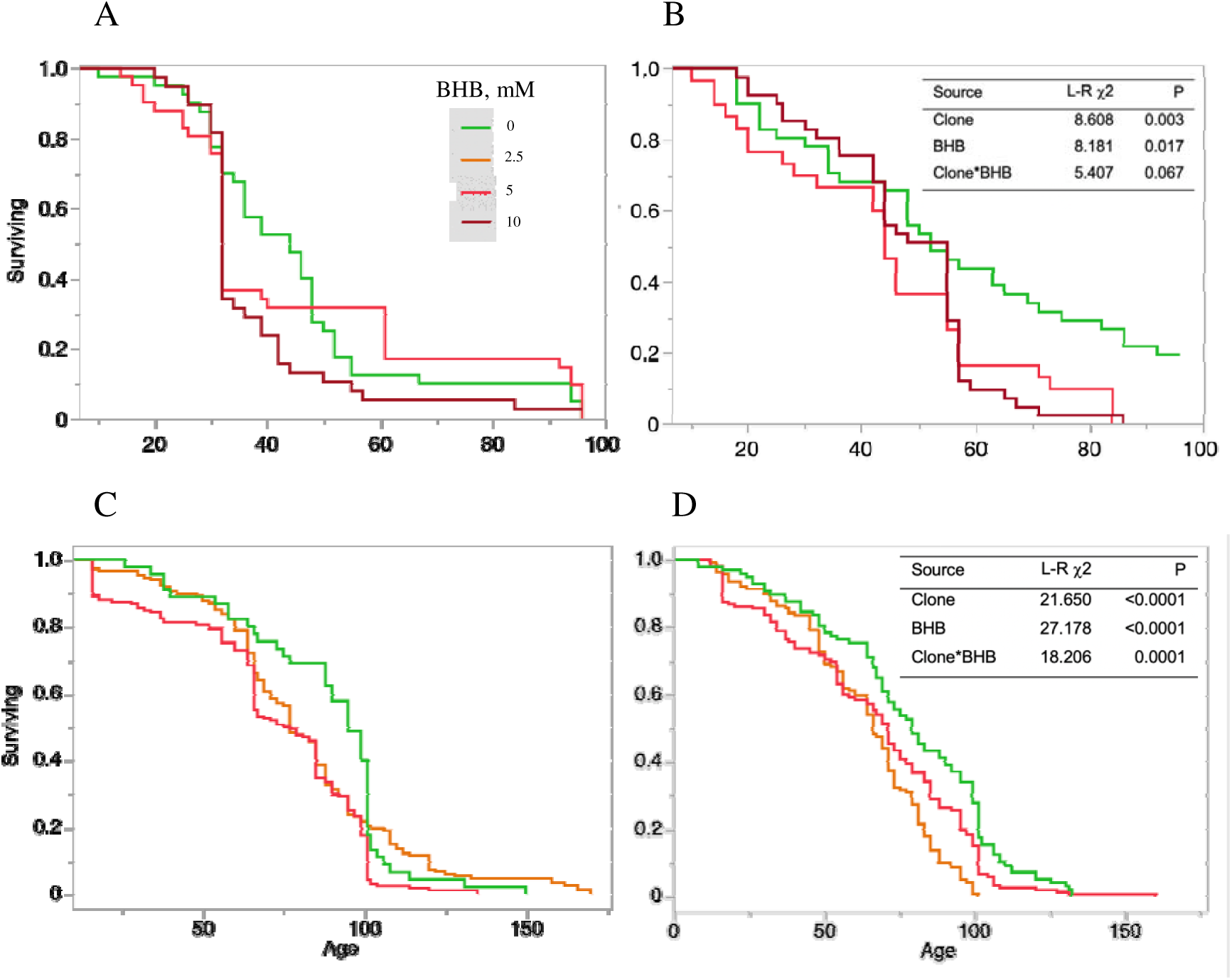
Lifetime BHB exposure reduces lifespan in *Daphnia* maintained at 1×10^5^ cells/mL/day at density of 1 individual per 20 mL, regardless of group size and experimental set up. A, B: experiment 1; C, D: experiment 2; A, C: clone GB-EL75-69; B, D: clone IL-M1-8. Inserts: Proportional hazards analysis of lifespan.

Intermittent BHB exposure of 20 days duration had no effect on survival past the onset of exposure in a combined analysis with all six cohorts with different ages of exposure onset analyzed together (Fig 2A). Median lifespan was higher in the BHB-exposed cohorts than in the control in all but the earliest exposure onset cohort (Fig. 2B), but the difference was not significant for any of the cohorts. See Supplementary Fig. S1 for the same data with each cohort shown separately.

**Fig. 2.**
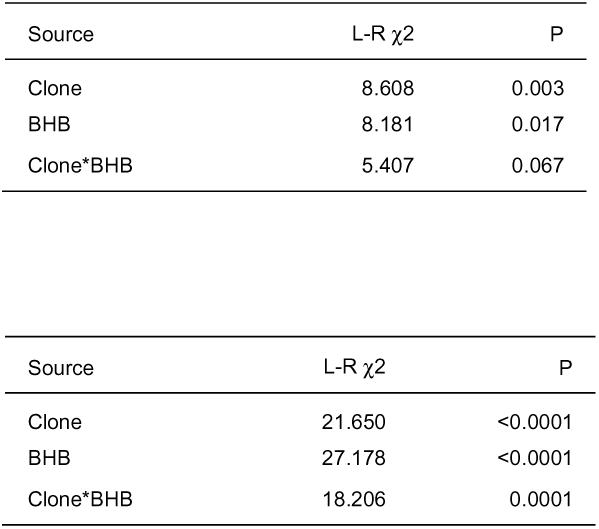
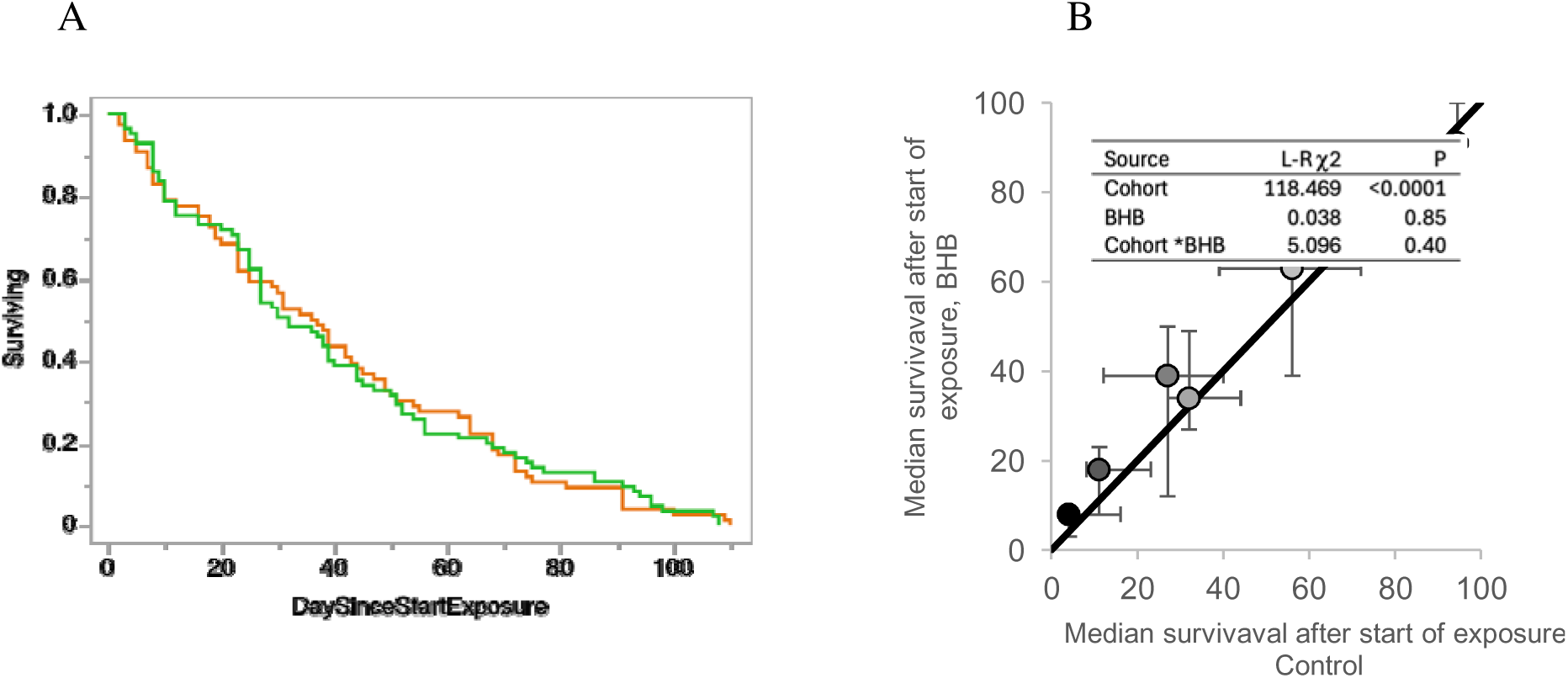
Short term exposure to 2.5 mM BHB does not affect post-exposure survival. A: survival curves (cohorts combined) in Experiment 3. Insert: Parametric survival test. B: comparison of median survival post exposure, BHB vs control, in 6 cohorts (symbols colored by the age of cohort at the start of exposure, from 15 to 144 days, white to black). Bars are 95% CIs.

### BHB exposure increased fecundity in food-limited Daphnia

Life-long continuous exposure to BHB increased *Daphnia* fecundity in both clones and nearly all age classes, with 5 mM exposure having a stronger effect that 10 mM exposure, where toxic effect was apparent, particularly in the oldest ages (Fig. 3; Table 2). In the oldest individuals this effect was comparable in size to that of two-fold food concentration (2 x10^5^ cells/mL/day), but it was much relatively weaker in younger individuals.

**Table 2.** Two-way REML ANOVA of fecundity differences between clones, life-long BHB treatments, and their interaction, with maternal age as a covariable, in Experiments 1 and 2. Replicate jars (Experiment 1) or tanks (Experiment 2) were included into the model and nested random variables nested within clones. Fecundity square root-transformed.

| Source | DF | DFDen | F Ratio | Prob > F |
| --- | --- | --- | --- | --- |
| Experiment 1: life-long exposure to 0, or 5, or 10 mM BHB,<br>group size 10 |  |  |  |  |
| BHB | 2 | 563.6 | 11.71 | <0.0001 |
| Clone | 1 | 23.1 | 0.55 | 0.47 |
| BHB*Clone | 2 | 548.3 | 2.57 | 0.08 |
| Age | 1 | 529.5 | 47.46 | <0.0001 |
| Experiment 2: life-long exposure to 0, or 2.5, or 5 mM BHB, life-long<br>exposure, group size 50. Ages 0 - 100 days |  |  |  |  |
| BHB | 2 | 435.5 | 5.00 | 0.0071 |
| Clone | 1 | 3.3 | 0.21 | 0.68 |
| BHB*Clone | 2 | 437.4 | 3.53 | 0.0300 |
| Age | 2 | 448.9 | 181.14 | <0.0001 |
| Experiment 2: life-long exposure to 0, or 2.5, or 5 mM BHB, life-long<br>exposure, group size 1. Ages 100 – 160 days |  |  |  |  |
| BHB | 2 | 168.2 | 2.23 | 0.11 |
| Clone | 1 | 2.9 | 0.34 | 0.60 |
| BHB*Clone | 2 | 163.4 | 0.33 | 0.72 |
| Age | 1 | 198.7 | 14.65 | 0.0002 |

Similarly, life-long intermittent (2 h/day) exposure to BHB at 2.5 and 5 mM concentrations (Experiment 2) caused increase in age-specific fecundity (clutch size). This increase also did not show a consistent dosage effect and also differed across age classes and between the two clones tested (Fig. 4, Table 1). Generally, the GB-EL75-69 clone responded to BHB exposure more consistently across age than the IL-M1-8 clone, but required a stronger concentration of BHB to respond (Fig. 4).

**Fig. 4.**
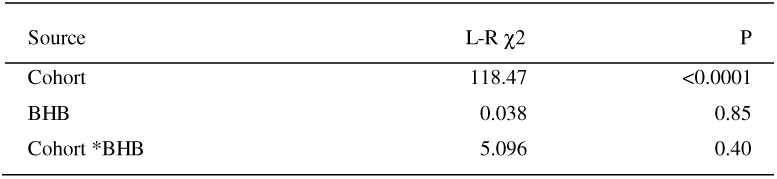
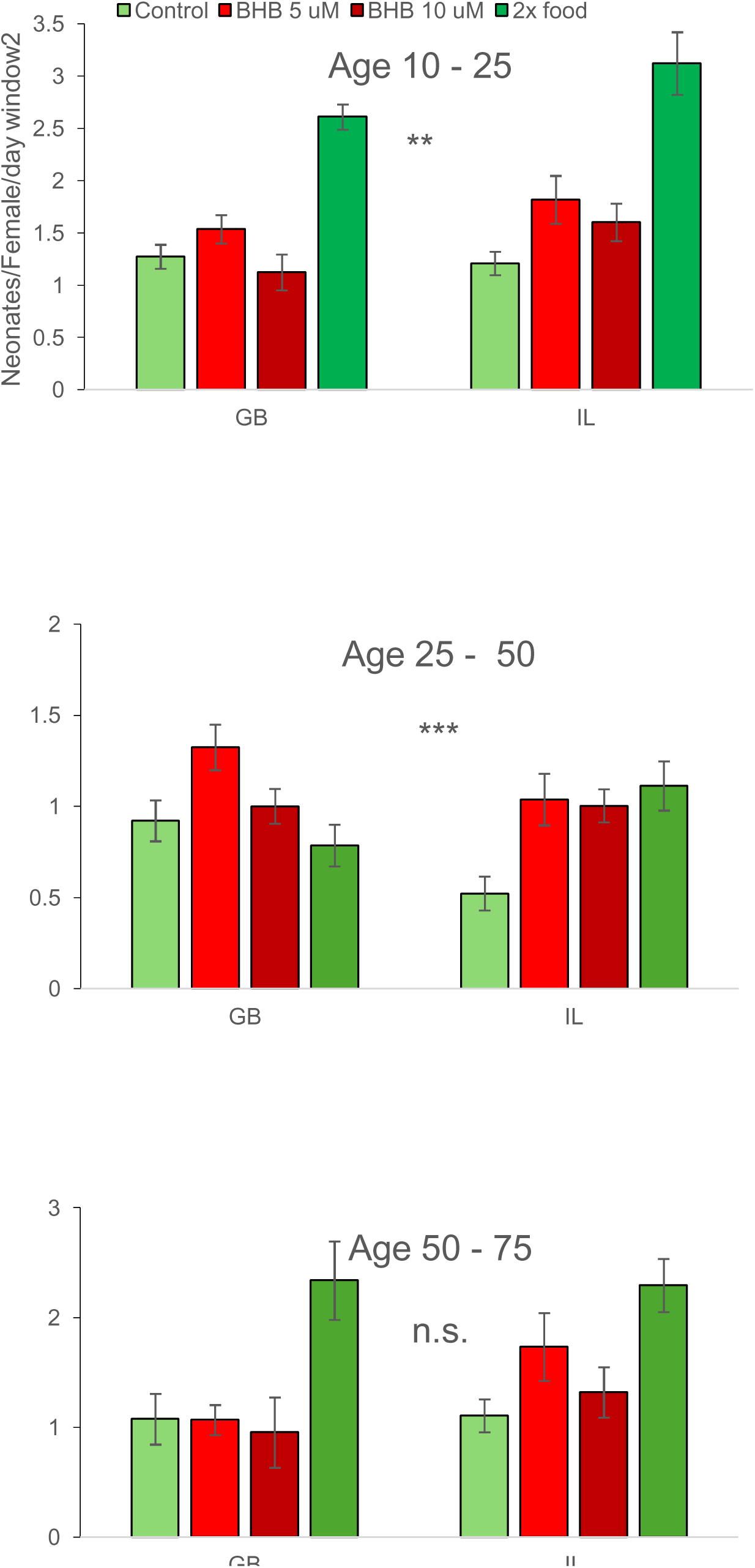
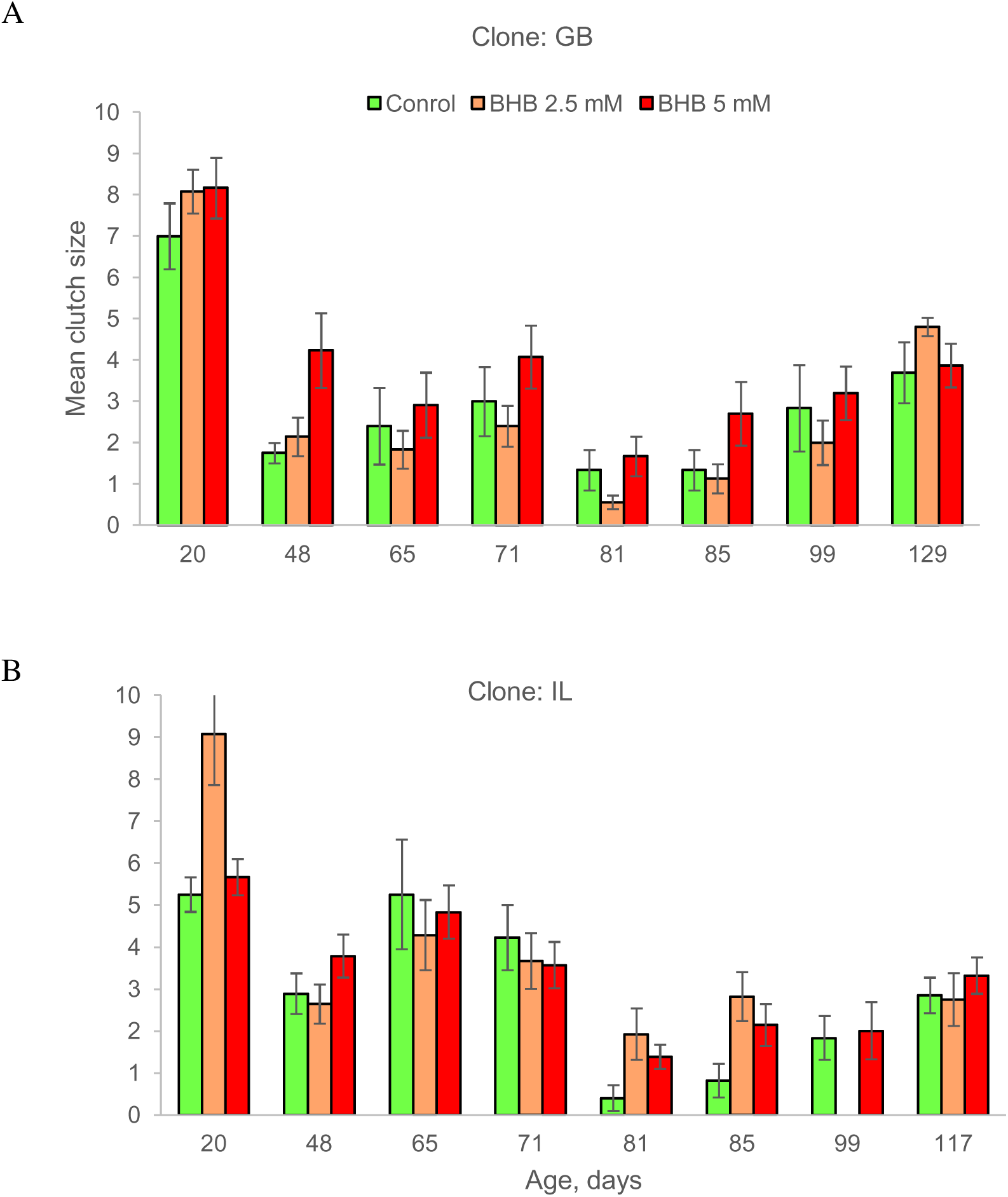
Age-specific fecundity (clutch size) in 2 *D. magna* clones (A: GB-EL75-69; B: IL-M1-8) after life-long intermittent exposure to either 2.5 mM (orange) or 5 mM BHB red vs. control treatment (green) in Experiment 2. See Table 2 for statistical analysis. All age classes except the oldest one represent clutches of females sampled from the experimental tanks; the oldest age class measured in individual females surviving to that age.

Fecundity of *Daphnia* exposed to 5 mM of BHB for 20-day periods at various ages responded differently in cohorts exposed at different ages (Fig. 5, Supplementary Table S2). In the cohort with a late onset of exposure (day 83 of age, Fig. 5A) the increase in fecundity was observed shortly (20-40 days) after the onset of exposure and the exposed daphnids remained reproductively active longer than the control ones (up to 160 days of age). The same late-age reproductive activity extending effect was observed in the cohort with the onset of exposure at the age of 70 days, although the effect of BHB was reversed shortly after exposure (Fig. 5B), with no significant effect of BHB overall. In the cohort in which BHB exposure started at the age of 40, the exposure increased fecundity only shortly after exposure (Fig. 5C). Finally, in the cohort with the earliest (day 15) onset of exposure, fecundity did not differ from that in the control cohort throughout the lifespan, but the late-life fecundity peak (Dua et al. 2024) was observed sooner than in the control cohort (Fig. 5D).

**Fig. 5.**
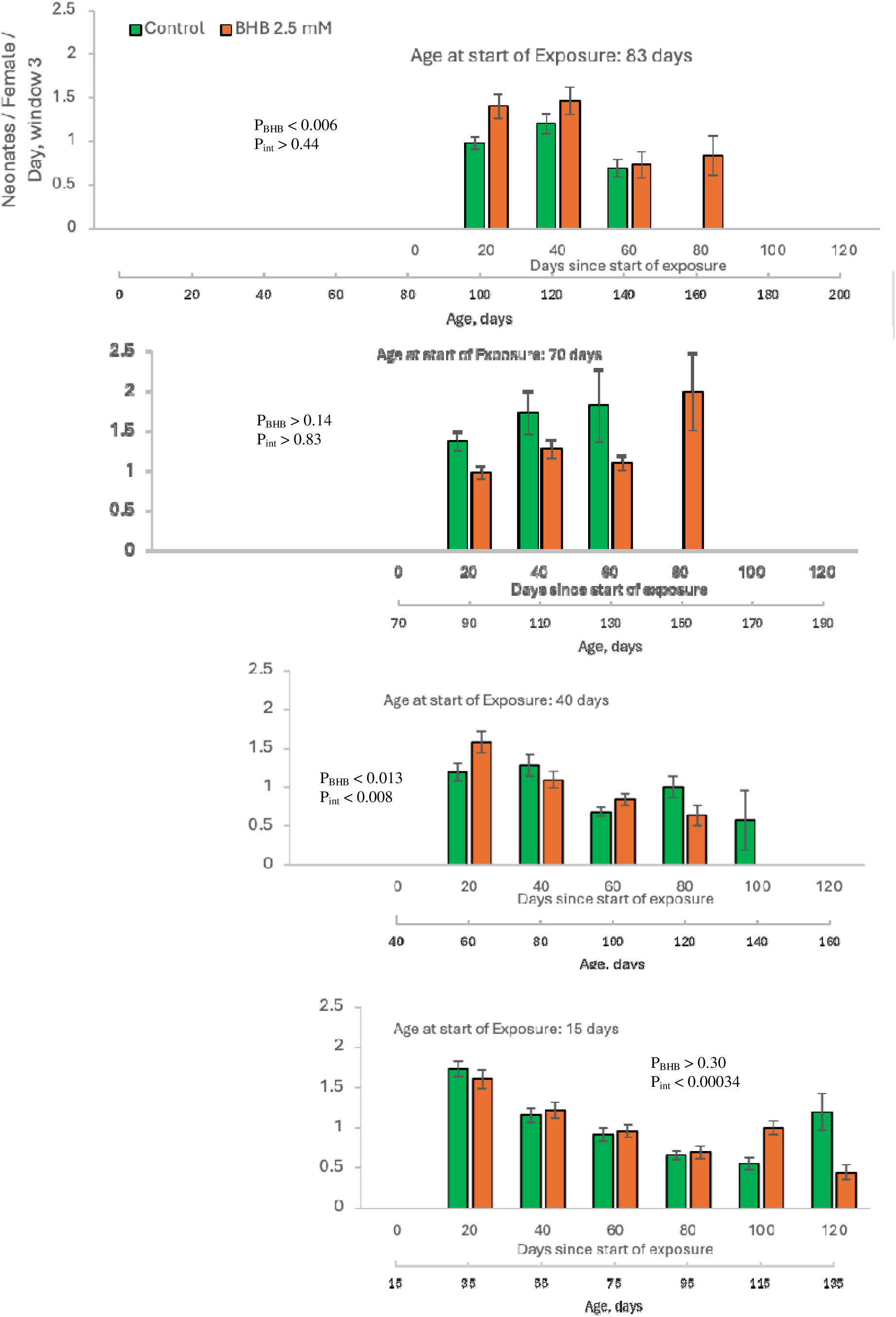
Fecundity in 4 cohorts of Experiment 3 (clone: IL) exposed to 2.5 mM BHB for 20 days at different ages, binned by 20 days since exposure start. Horizonal axis: days since the start of exposure and astronomical age. Bars are SEs. Inserts indicate the results of a 2-way ANOVA with binned days since exposure start and BHB vs. control treatment as factors; top P-value for the BHB effect, bottom for the interaction effect (see Supplementary Table S1 for full statistical results).

### Transgenerational effects of maternal exposure to 2.5 mM BHB

We observed mild maternal and transgenerational effects of HBH exposure on life history of the offspring of females intermittently exposed to BHB relative to control, although these effects are highly genotype-specific (Table 3). First, in 1 out of 3 clones tested, BHB-exposed mothers produced smaller (Fig. 6A) but better lipid-provisioned (Fig. 6B) offspring. Neonates’ body size, at the same time, was not correlated with lipid content when differences among clones or between treatments are accounted for (Supplementary Table S2). In two other clones, sisters of neonates in which body length and lipid content at birth were measured were, on the other hand, larger at maturity (at the age of production of the first clutch; Fig. 6C). Finally, two out of three clones tested showed the opposite effects of maternal exposure to BHB in terms of offspring number in the first clutch (Fig. 6D), as reflected by a significant maternal BHB x clone interaction terms (Table 3). In no further clutches there were any differences in offspring number between daughters of BHB-exposed and control mothers (data not reported); nor there were any differences in longevity between these two groups (Fig. 6E).

**Fig. 6.**
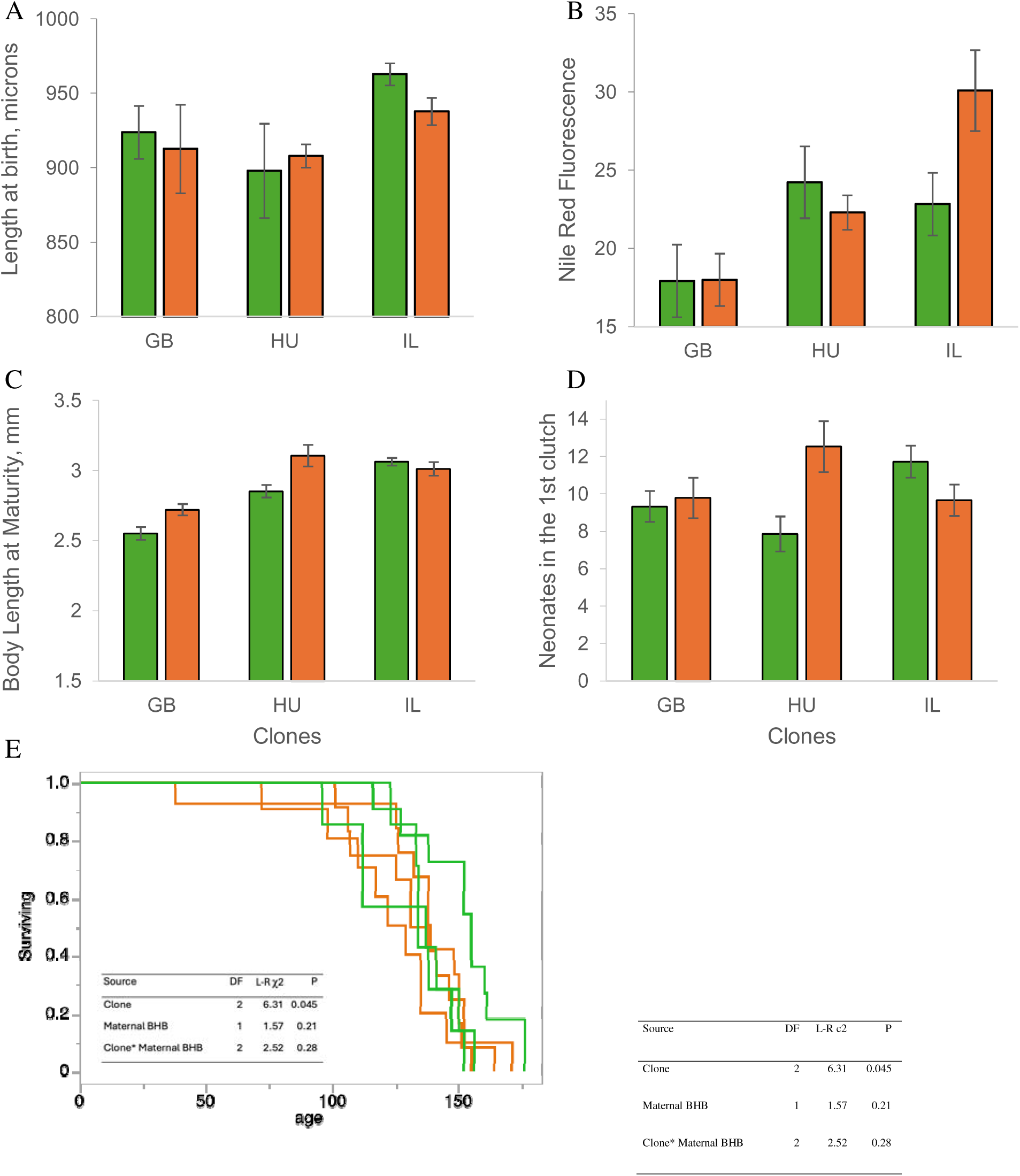
Body size (A), lipid content (Nile Red fluorescence, arbitrary units, B), size at maturity (C), fecundity (size of the 1^st^ clutch, D), and longevity (E; insert: Proportional Hazards test) of offspring produced by females exposed to either 2.5 mM BHB or ADaM medium (control) for 20 days prior to their birth. See Table 3 for statistical analysis.

**Table 3.** Two-way REML ANOVA of body size and lipid content (median Nile Red fluorescence, background-subtracted) at birth, body length at maturity, and number of offspring in the 1^st^ clutch in offspring of females exposed to either 2.5 mM BHB or ADaM medium (control) for 20 days prior to their birth (Fig. 5 A, B). Maternal ID used as a random effect nested within Clones. See Supplementary Table S2 for joint analysis of length at birth and lipid content.

| Source | DF | DFDen | F Ratio | Prob > F |
| --- | --- | --- | --- | --- |
| Response: Length at birth |  |  |  |  |
| Maternal BHB | 1 | 51.7 | 0.677 | 0.41 |
| Clone | 2 | 50.7 | 10.226 | 0.0002 |
| Clone*Maternal BHB | 2 | 50.7 | 1.479 | 0.24 |
| Response: Nile Red fluorescence |  |  |  |  |
| Maternal BHB | 1 | 41.5 | 1.55 | 0.22 |
| Clone | 2 | 41 | 10.99 | 0.0002 |
| Clone*Maternal BHB | 2 | 41 | 4.33 | 0.02 |
| Response: Length at maturity |  |  |  |  |
| Clone | 2 | 9.936 | 23.24 | 0.0002 |
| Maternal BHB | 1 | 343.4 | 59.09 | <0.0001 |
| Clone*Maternal BHB | 2 | 342.3 | 21.61 | <0.0001 |
| Response: Neonates, 1<sup>st</sup> clutch |  |  |  |  |
| Clone | 2 | 6.622 | 0.45 | 0.65 |
| Maternal BHB | 1 | 70.74 | 1.30 | 0.26 |
| Clone*Maternal BHB | 2 | 69.84 | 4.99 | 0.01 |

## Discussion

We observed decreased lifespan in food-limited *Daphnia* continuously or intermittently exposed to 2.5 – 10 mM of betahydroxybutyrate for life. No change in life expectancy was observed after 20-day exposure this compound. In a separate study (Pearson and Yampolsky, submitted), we, on the other hand, observed decrease of early-life mortality under intermittent exposure to BHB in *Daphnia* fed *ad libitum* (4 times the food availability in experiments reported here), matching the decrease in mortality induced by food limitation. This indicates that BHB does not affect lifespan *per se,* but rather mimics caloric restriction effect, shortening life of dietary limited, but extending life of overfed *Daphnia.* On the other hand, BHB exposure induced higher fecundity, in some cases nearly doubling the number of offspring produced. Although the magnitude of this effect differed greatly between the two reference clones tested and between age classes, as reflected by appropriate significant interaction terms, there is little doubt that BHB supplementation increased investment into reproduction in food-limited *Daphnia*. Furthermore, this effect is transgenerational, at least for 1 out of three clones tested (limited to the increase in the size of the very first clutch of the daughters of BHB-exposed mothers).

Two related questions arise with the interpretation of these facts. The first question is whether the external BHB source was utilized as an additional resource allowing increased egg production, or a signal to do so. The second question is whether the transgenerational effect occurs through, again, differences in provisioning of offspring with extra resources, or through sending them a signal.

A compelling argument can be set forth favoring the signaling that then energetic mechanism of the observed increase in fecundity. It is fully implausible that the BHB concentrations at which fecundity increases are the most apparent (2.5 – 5 mM exposure solution) may be sufficient to supply *Daphnia* with enough extra ATP to save resources for increased investments into egg production. Indeed, if one assumes that during a 2-hour exposure daphnids achieve internal concentration of BHB at diffusion equilibrium with the external concentration, then the 5 mM exposure regimen would result in 10 nmoles of BHB consumed. When fully oxidized this amount of BHB would generate the amount of ATP similar to that generated by full oxidation of 6.7 nmoles of glucose, and, if the two catabolic pathways are equivalent in terms of biomass consumed, this would result in savings of about 1.2 ug of biomass per day. This is equivalent to about 0.1 extra egg produced per day or less than half an egg per clutch (assuming 0.1 ug dry weight of a *D. magna* egg, Trubetskova & Lampert 1995). Certainly more efficient consumption of BHB than simple diffusion without immediate utilization would, in principle, result in higher total amount of BHB consumed and a greater biomass gain, but it is still difficult to envision this to acount for the observed fecundity. Furthermore, direct energy supply explanation is incompatible with a long-term consequences of early-or mid-life 20-day exposure to BHB (Fig. 5) and, in particular, with a transgenerational fecundity increasing effect observed in the HU-K-6 clone (Fig. 6D). Rather, one should assume that one of numerous signaling functions of BHB is at play, perhaps those affecting lipid metabolism or NAD+/NADP ratio (Newman and Verdin 2017).

Similarly, it is unlikely that the genotype-specific transgenerational increase of early fecundity in daughters of BHB exposed mothers is unlikely to occur through extra provisioning of germline cells. First, the daughter generation was exposed to BHB in the form of oocyte precursor cells, one ovary cycle before oocyte provisioning starts. Second, the clone showing the transgenerational increase in growth rate to maturity and early fecundity (HU clone; Fig. 6 C,D) showed no changes in body size or lipid content at birth (Fig. 6A, B), in fact showing slightly lower lipid content at birth. Conversely, the clone which did show a significant increase in lipid provisioning by BHB-exposed mothers (IL clone, Fig. 6B), possibly with a trade-off with neonate body size (Fig. 6A), showed reduced, not increased early mortality (Fig. 6D). Thus thre is no evidence that transgenerational effect is based on biomass or other resources allocations into offspring.

Given the fact that organisms as diverse as *Daphnia* (Tessier & Goulden 1982; Martin-Creuzburg et al. 2009) and cattle (Missio et al. 2022) reproductive functions depend on utilization of body fat reserves, BHB regulatory functions affecting nutrient availability signaling and lipid metabolism may be a mechanism of the observed fecundity changes. Thus, the significant differences among genotypes in how these signals are transmitted within and across generations are particularly intriguing. A broader survey of *Daphnia* genotypes with respect to their response to extraneous BHB supplementation followed by a genome-wide analysis might uncover genes responsible for lipid metabolism and reproductive allocation in this model organism.

## Acknowledgements

We are grateful to Patrick Bradshaw for useful discussions and to members of Yampolsky lab for laboratory assistance. This work was supported by Impetus Foundation grant to LYY.

## Notes

### Competing Interest Statement

The authors have declared no competing interest.

## References

1. Agrelius TC, Altman J, Dudycha JL. The maternal effects of dietary restriction on Dnmt expression and reproduction in two clones of Daphnia pulex. Heredity (Edinb). 2023 Feb;130(2):73–81. doi: 10.1038/s41437-022-00581-7. Epub 2022 Dec 7. PMID: 36477021; PMCID: PMC9905607.

2. Agrelius TC, Dudycha JL. Maternal effects in the model system Daphnia: the ecological past meets the epigenetic future. Heredity (Edinb). 2025 Jan 8. doi: 10.1038/s41437-024-00742-w. Epub ahead of print. PMID: 39779907.

3. Anderson CE, Malek MC, Jonas-Closs RA, Cho Y, Peshkin L, Kirschner MW, Yampolsky LY. Inverse Lansing effect: maternal age and provisioning affecting daughters’ longevity and male offspring production. Am Nat. 2022 Nov;200(5):704–721. doi: 10.1086/721148. Epub 2022 Sep 28. PMID: 36260845.

4. Andrewartha SJ, Burggren WW. Transgenerational variation in metabolism and life-history traits induced by maternal hypoxia in Daphnia magna. Physiol Biochem Zool. 2012 Nov-Dec;85(6):625–34. doi: 10.1086/666657. Epub 2012 Aug 2. PMID: 23099460.

5. Balietti M, Casoli T, Di Stefano G, Giorgetti B, Aicardi G, Fattoretti P. Ketogenic diets: an historical antiepileptic therapy with promising potentialities for the aging brain. Ageing Res Rev. 2010 Jul;9(3):273–9. doi: 10.1016/j.arr.2010.02.003. Epub 2010 Feb 24. PMID: 20188215.

6. Ben-Ami F, Ebert D, Regoes RR. Pathogen dose infectivity curves as a method to analyze the distribution of host susceptibility: a quantitative assessment of maternal effects after food stress and pathogen exposure. Am Nat. 2010 Jan;175(1):106–15. doi: 10.1086/648672. PMID: 19911987.

7. Boersma M. 1995. The allocation of resources to reproduction in Daphnia galeata: against the odds? Ecology 76(4):1251–1261.

8. Boersma M. 1997. Offspring size in Daphnia: does it pay to be overweight? Hydrobiologia 360: 79–88, 1997.

9. Campisi J, Kapahi P, Lithgow GJ, Melov S, Newman JC, Verdin E. From discoveries in ageing research to therapeutics for healthy ageing. Nature. 2019 Jul;571(7764):183–192. doi: 10.1038/s41586-019-1365-2. Epub 2019 Jul 10. PMID: 31292558; PMCID: PMC7205183.

10. Cho, Y., R.A. Jonas-Closs, L.Y. Yampolsky, M.W. Kirschner, L. Peshkin. 2022. “Smart Tanks”: An Intelligent High-Throughput Intervention Testing Platform in *Daphnia*. Aging Cell 2022: e13571. doi: 10.1111/acel.13571

11. Coakley CM, Nestoros E, Little TJ. Testing hypotheses for maternal effects in Daphnia magna. J Evol Biol. 2018 Feb;31(2):211–216. doi: 10.1111/jeb.13206. Epub 2017 Nov 22. PMID: 29117456; PMCID: PMC6849578.

12. Dao TS, Vo TM, Wiegand C, Bui BT, Dinh KV. Transgenerational effects of cyanobacterial toxins on a tropical micro-crustacean Daphnia lumholtzi across three generations. Environ Pollut. 2018 Dec;243(Pt B):791–799. doi: 10.1016/j.envpol.2018.09.055. Epub 2018 Sep 15. PMID: 30241003.

13. Edwards C, Canfield J, Copes N, Rehan M, Lipps D, Bradshaw PC. D-beta-hydroxybutyrate extends lifespan in C. elegans. Aging (Albany NY). 2014 Aug;6(8):621–44. doi: 10.18632/aging.100683. PMID: 25127866; PMCID: PMC4169858.

14. Feiner N, Radersma R, Vasquez L, Ringnér M, Nystedt B, Raine A, Tobi EW, Heijmans BT, Uller T. Environmentally induced DNA methylation is inherited across generations in an aquatic keystone species. iScience. 2022 Apr 25;25(5):104303. doi: 10.1016/j.isci.2022.104303. PMID: 35573201; PMCID: PMC9097707.

15. Frost PC, Ebert D, Larson JH, Marcus MA, Wagner ND, Zalewski A. Transgenerational effects of poor elemental food quality on Daphnia magna. Oecologia. 2010 Apr;162(4):865–72. doi: 10.1007/s00442-009-1517-4. Epub 2009 Dec 2. PMID: 19957090.

16. Garbutt JS, Little TJ. Maternal food quantity affects offspring feeding rate in Daphnia magna. Biol Lett. 2014 Jul;10(7):20140356. doi: 10.1098/rsbl.2014.0356. PMID: 25030044; PMCID: PMC4126628.

17. Garbutt JS, Little TJ. Bigger is better: changes in body size explain a maternal effect of food on offspring disease resistance. Ecol Evol. 2017 Feb 3;7(5):1403–1409. doi: 10.1002/ece3.2709. PMID: 28261452; PMCID: PMC5330872.

18. Garbutt JS, Scholefield JA, Vale PF, Little TJ. Elevated maternal temperature enhances offspring disease resistance in Daphnia magna. Funct Ecol 2014; 28: 424–431.

19. Ghosh HS. 2008 The anti-aging, metabolism potential of SIRT1. Curr Opin Investig Drugs 9(10): 1095–1102. PMID: 18821472.

20. Gliwicz ZM, Guisande C. Family planning in Daphnia: resistance to starvation in offspring born to mothers grown at different food levels. Oecologia. 1992 Oct;91(4):463–467. doi: 10.1007/BF00650317. PMID: 28313496.

21. Glozak MA, Sengupta N, Zhang X, Seto E. 2005. Acetylation and deacetylation of non-histone proteins. Gene 363:15–23)

22. Guisande, C., & Gliwicz, Z. M. (1992). Egg size and clutch size in two Daphnia species grown at different food levels. Journal of Plankton Research, 14(7), 997–1007. doi:10.1093/plankt/14.7.997

23. Han YM, Ramprasath T, Zou MH. β-hydroxybutyrate and its metabolic effects on age-associated pathology. Exp Mol Med. 2020 Apr;52(4):548–555. doi: 10.1038/s12276-020-0415-z. Epub 2020 Apr 8. PMID: 32269287; PMCID: PMC7210293.

24. Harney E, Paterson S, Collin H, Chan BHK, Bennett D, Plaistow SJ. Pollution induces epigenetic effects that are stably transmitted across multiple generations. Evol Lett. 2022 Feb 3;6(2):118–135. doi: 10.1002/evl3.273. PMID: 35386832; PMCID: PMC8966472.

25. Hearn J, Chow FW, Barton H, Tung M, Wilson PJ, Blaxter M, Buck A, Little TJ. *Daphnia* magna microRNAs respond to nutritional stress and ageing but are not transgenerational. Mol Ecol. 2018 Mar;27(6):1402–1412. doi: 10.1111/mec.14525. Epub 2018 Mar 7. PMID: 29420841.

26. Im H, Kang J, Jacob MF, Bae H, Oh JE. Transgenerational effects of benzotriazole on the gene expression, growth, and reproduction of Daphnia magna. Environ Pollut. 2023 Apr 15;323:121211. doi: 10.1016/j.envpol.2023.121211. Epub 2023 Feb 3. PMID: 36740167.

27. Jeremias G, Barbosa J, Marques SM, De Schamphelaere KAC, Van Nieuwerburgh F, Deforce D, Gonçalves FJM, Pereira JL, Asselman J. Transgenerational Inheritance of DNA Hypomethylation in Daphnia magna in Response to Salinity Stress. Environ Sci Technol. 2018 Sep 4;52(17):10114–10123. doi: 10.1021/acs.est.8b03225. Epub 2018 Aug 24. PMID: 30113818.

28. Jiang X, Yang W, Zhao S, Liang H, Zhao Y, Chen L, Li R. Maternal effects of inducible tolerance against the toxic cyanobacterium Microcystis aeruginosa in the grazer Daphnia carinata. Environ Pollut. 2013 Jul;178:142–6. doi: 10.1016/j.envpol.2013.03.017. Epub 2013 Apr 9. PMID: 23570781.

29. Kenyon CJ. 2010. The genetics of ageing. Nature 464:504–512

30. Klüttgen B, U Dülmer, M Engels, H.T Ratte. 1994. AdaM, an artificial freshwater for the culture of zooplankton. Water Research 28 (3), 743–746.10.1016/0043-1354(94)90157-0.

31. Kulak D, Polotsky AJ. Should the ketogenic diet be considered for enhancing fertility? Maturitas. 2013 Jan;74(1):10–3. doi: 10.1016/j.maturitas.2012.10.003. Epub 2012 Oct 31. PMID: 23122539.

32. Kwon HS, Ott M. 2008. The ups and downs of SIRT1. Trends Biochem Sci 33(11): 517–525. doi: 10.1016/j.tibs.2008.08.001. PMID: 18805010.

33. LaMontagne, J.M. and McCauley, E. (2001), Maternal effects in Daphnia: what mothers are telling their offspring and do they listen?. Ecology Letters, 4: 64–71. 10.1046/j.1461-0248.2001.00197.x

34. Lai KP, Li JW, Chan CY, Chan TF, Yuen KW, Chiu JM. Transcriptomic alterations in Daphnia magna embryos from mothers exposed to hypoxia. Aquat Toxicol. 2016 Aug;177:454–63. doi: 10.1016/j.aquatox.2016.06.020. Epub 2016 Jun 24. PMID: 27399157.

35. LeBlanc GA, Wang YH, Holmes CN, Kwon G, Medlock EK. A transgenerational endocrine signaling pathway in Crustacea. PLoS One. 2013 Apr 17;8(4):e61715. doi: 10.1371/journal.pone.0061715. PMID: 23613913; PMCID: PMC3629115.

36. Lee JS, Oh Y, Lee JS, Kim HS. Acute toxicity, oxidative stress, and apoptosis due to short-term triclosan exposure and multi– and transgenerational effects on in vivo endpoints, antioxidant defense, and DNA damage response in the freshwater water flea Daphnia magna. Sci Total Environ. 2023 Mar 15;864:160925. doi: 10.1016/j.scitotenv.2022.160925. Epub 2022 Dec 18. PMID: 36543274.

37. Lyu K, Zhang L, Gu L, Zhu X, Wilson AE, Yang Z. Cladoceran offspring tolerance to toxic Microcystis is promoted by maternal warming. Environ Pollut. 2017 Aug;227:451–459. doi: 10.1016/j.envpol.2017.04.095. Epub 2017 May 6. PMID: 28486188.

38. Martin-Creuzburg D, Sperfeld E, Wacker A. 2009. Colimitation of a freshwater herbivore by sterols and polyunsaturated fatty acids. Proc Biol Sci. 2009 May 22;276(1663):1805–14. doi: 10.1098/rspb.2008.1540. Epub 2009 Feb 20. PMID: 19324803; PMCID: PMC2674483.

39. Madhavan SS, Stubbs BJ. Beta-hydroxybutyrate. Trends Endocrinol Metab. 2025 Jan;36(1):96-97. doi: 10.1016/j.tem.2024.06.005. Epub 2024 Jul 17. PMID: 39765208; PMCID: PMC11707391.

40. Martins A, Guilhermino L. Transgenerational effects and recovery of microplastics exposure in model populations of the freshwater cladoceran Daphnia magna Straus. Sci Total Environ. 2018 Aug 1;631-632:421–428. doi: 10.1016/j.scitotenv.2018.03.054. Epub 2018 Mar 16. PMID: 29529430.

41. Matsuyama S, Kimura K. Regulation of gonadotropin secretion by monitoring energy availability. Reprod Med Biol. 2014 Sep 24;14(2):39–47. doi: 10.1007/s12522-014-0194-0. PMID: 29259401.

42. Mckee D, Ebert D. The interactive effects of temperature, food level and maternal phenotype on offspring size in Daphnia magna. Oecologia. 1996 Jul;107(2):189–196. doi: 10.1007/BF00327902. PMID: 28307304.

43. Michalaki A, Grintzalis K. Acute and transgenerational effects of non-steroidal anti-inflammatory drugs on Daphnia magna. Toxics. 2023 Mar 29;11(4):320. doi: 10.3390/toxics11040320. PMID: 37112547; PMCID: PMC10145367.

44. Michel J, Ebert D, Hall MD. The trans-generational impact of population density signals on host-parasite interactions. BMC Evol Biol. 2016 Nov 25;16(1):254. doi: 10.1186/s12862-016-0828-4. PMID: 27887563; PMCID: PMC5123254.

45. Mikulski A, Mazurczak D. Maternal effect in salinity tolerance of Daphnia-One species, various patterns? PLoS One. 2023 Apr 4;18(4):e0283546. doi: 10.1371/journal.pone.0283546. PMID: 37014884; PMCID: PMC10072381.

46. Missio D, Fritzen A, Cupper Vieira C, Germano Ferst J, Farias Fiorenza M, Guedes de Andrade L, Martins de Menezes B, Bisneto, Tomazele Rovani M, Gazieira Gasperin B, Dias Gonçalves PB, Ferreira R. 2022. Increased β-hydroxybutyrate (BHBA) concentration affect follicular growth in cattle. Anim Reprod Sci. 2022 Aug;243:107033. doi: 10.1016/j.anireprosci.2022.107033. Epub 2022 Jul 6. PMID: 35816934.

47. Møller N. 2020. Ketone Body, 3-Hydroxybutyrate: Minor Metabolite – Major Medical Manifestations. J Clin Endocrinol Metab. 2020 Sep 1;105(9):dgaa370. doi: 10.1210/clinem/dgaa370. PMID: 32525972.

48. Newman JC, Covarrubias AJ, Zhao M, Yu X, Gut P, Ng CP, Huang Y, Haldar S, Verdin E. Ketogenic Diet Reduces Midlife Mortality and Improves Memory in Aging Mice. Cell Metab. 2017 Sep 5;26(3):547–557.e8. doi: 10.1016/j.cmet.2017.08.004. PMID: 28877458; PMCID: PMC5605815.

49. Newman JC & Verdin E. 2014. Ketone bodies as signaling metabolites. Trends in Endocrinology and Metabolism, 25(1), 42–52. 10.1016/j.tem.2013.09.002

50. Newman JC, Verdin E. 2017. β-Hydroxybutyrate: A Signaling Metabolite. Annu Rev Nutr. 2017 Aug 21;37:51–76. doi: 10.1146/annurev-nutr-071816-064916. PMID: 28826372; PMCID: PMC6640868.

51. Padilla Suarez EG, Pugliese S, Galdiero E, Guida M, Libralato G, Saviano L, Spampinato M, Pappalardo C, Siciliano A. Multigenerational tests on Daphnia spp.: a vision and new perspectives. Environ Pollut. 2023 Nov 15;337:122629. doi: 10.1016/j.envpol.2023.122629. Epub 2023 Sep 27. PMID: 37775025.

52. Paraskevopoulou S, Gattis S, Ben-Ami F. Parasite resistance and parasite tolerance: insights into transgenerational immune priming in an invertebrate host. Biol Lett. 2022 Apr;18(4):20220018. doi: 10.1098/rsbl.2022.0018. Epub 2022 Apr 6. PMID: 35382587; PMCID: PMC8984330.

53. Plaistow SJ, Shirley C, Collin H, Cornell SJ, Harney ED. Offspring Provisioning Explains Clone-Specific Maternal Age Effects on Life History and Life Span in the Water Flea, Daphnia pulex. Am Nat. 2015 Sep;186(3):376–89. doi: 10.1086/682277. Epub 2015 Jul 2. PMID: 26655355.

54. Qi Q, Li Q, Li J, Mo J, Tian Y, Guo J. Transcriptomic analysis and transgenerational effects of ZnO nanoparticles on Daphnia magna: Endocrine-disrupting potential and energy metabolism. Chemosphere. 2022 Mar;290:133362. doi: 10.1016/j.chemosphere.2021.133362. Epub 2021 Dec 18. PMID: 34933032.

55. Radersma R, Hegg A, Noble DWA, Uller T. Timing of maternal exposure to toxic cyanobacteria and offspring fitness in Daphnia magna: Implications for the evolution of anticipatory maternal effects. Ecol Evol. 2018 Nov 20;8(24):12727–12736. doi: 10.1002/ece3.4700. PMID: 30619577; PMCID: PMC6309005.

56. Sakwińska O.2003. Persistent Maternal Identity Effects on Life History Traits in Daphnia. Oecologia 138, No. 3, pp. 379–386.

57. Sarapultseva EI, Dubrova YE. The long-term effects of acute exposure to ionising radiation on survival and fertility in Daphnia magna. Environ Res. 2016 Oct;150:138–143. doi: 10.1016/j.envres.2016.05.046. Epub 2016 Jun 8. PMID: 27288911.

58. Sha Y, Tesson SVM, Hansson LA. Diverging responses to threats across generations in zooplankton. Ecology. 2020 Nov;101(11):e03145. doi: 10.1002/ecy.3145. Epub 2020 Aug 19. PMID: 32740928; PMCID: PMC7685145.

59. Shahmohamadloo RS, Fryxell JM, Rudman SM. Transgenerational epigenetic inheritance increases trait variation but is not adaptive. bioRxiv [Preprint]. 2024 Apr 20:2024.04.15.589575. doi: 10.1101/2024.04.15.589575. PMID: 38659883; PMCID: PMC11042258.

60. Shimazu T, Hirschey MD, Newman J, He W, Shirakawa K, et al. 2013. Suppression of oxidative stress by β-hydroxybutyrate, an endogenous histone deacetylase inhibitor. Science 339: 211–214

61. Stein, L.R. and Imai, S. (2012) The dynamic regulation of NAD metabolism in mitochondria. Trends Endocrinol. Metab. 23: 420–428.

62. Stjernman M, Little TJ. 2011. Genetic variation for maternal effects on parasite susceptibility. J Evol Biol. 2011 Nov;24(11):2357–63. doi: 10.1111/j.1420-9101.2011.02363.x. Epub 2011 Aug 16. PMID: 21848987.

63. Sugishita Y, Suzuki-Takahashi Y, Yudoh K. 2024. Nicotinamide Adenine Dinucleotide (NAD)-Dependent Protein Deacetylase, Sirtuin, as a Biomarker of Healthy Life Expectancy: A Mini-Review. Curr Aging Sci. doi: 10.2174/0118746098319674240827104612. Epub ahead of print. PMID: 39377386.

64. Sun SJ, Dziuba MK, Jaye RN, Duffy MA. Transgenerational plasticity in a zooplankton in response to elevated temperature and parasitism. Ecol Evol. 2023 Feb 3;13(2):e9767. doi: 10.1002/ece3.9767. PMID: 36760704; PMCID: PMC9897957.

65. Tessier AJ, Goulden CE. 1982. Estimating food limitation in cladoceran populations, Limnology and Oceanography 27: 707–717. doi: 10.4319/lo.1982.27.4.0707.

66. Toyota K, Cambronero Cuenca M, Dhandapani V, Suppa A, Rossi V, Colbourne JK, Orsini L. Transgenerational response to early spring warming in Daphnia. Sci Rep. 2019 Mar 14;9(1):4449. doi: 10.1038/s41598-019-40946-3. PMID: 30872717; PMCID: PMC6418131.

67. Trubetskova, I., Lampert, W. 1995. Egg size and egg mass of Daphnia magna: response to food availability. In: Larsson, P., Weider, L.J. (eds) Cladocera as Model Organisms in Biology. Developments in Hydrobiology, vol 107. Springer, Dordrecht. 10.1007/978-94-011-0021-2_15

68. Vandegehuchte MB, Lemière F, Vanhaecke L, Vanden Berghe W, Janssen CR. Direct and transgenerational impact on Daphnia magna of chemicals with a known effect on DNA methylation. Comp Biochem Physiol C Toxicol Pharmacol. 2010 Apr;151(3):278–85. doi: 10.1016/j.cbpc.2009.11.007. Epub 2009 Dec 2. PMID: 19961956.

69. Veech RL. 2004. The therapeutic implications of ketone bodies: the effects of ketone bodies in pathological conditions: ketosis, ketogenic diet, redox states, insulin resistance, and mitochondrial metabolism. Prostaglandins Leukot. Essent. Fatty Acids 70:309–19

70. Veech RL, Bradshaw PC, Clarke K, Curtis W, Pawlosky R, King MT. Ketone bodies mimic the life span extending properties of caloric restriction. IUBMB Life. 2017 May;69(5):305–314. doi: 10.1002/iub.1627. Epub 2017 Apr 3. PMID: 28371201.

71. Wade GN, Schneider JE. Metabolic fuels and reproduction in female mammals. Neurosci Biobehav Rev. 1992 Summer;16(2):235–72. doi: 10.1016/s0149-7634(05)80183-6. PMID: 1630733.

72. Walsh MR, Castoe TA, Holmes J, Packer M, Biles K, Walsh MJ, Munch SB, Post DM (2016). Local adaptation in transgenerational responses to prey. Proceedings of the Royal Society of London B: Biological Sciences, 283: 20152271.

73. Walsh MR, Cooley F 4th, Biles K, Munch SB. Predator-induced phenotypic plasticity within– and across-generations: a challenge for theory? Proc Biol Sci. 2015 Jan 7;282(1798):20142205. doi: 10.1098/rspb.2014.2205. PMID: 25392477; PMCID: PMC4262177.

74. Walsh MR, Gillis MK. Transgenerational plasticity in the eye size of Daphnia. Biol Lett. 2021 Jun;17(6):20210143. doi: 10.1098/rsbl.2021.0143. Epub 2021 Jun 16. PMID: 34129799; PMCID: PMC8205523.

75. Walsh MR, Whittington D, Funkhouser C. Thermal transgenerational plasticity in natural populations of Daphnia. Integr Comp Biol. 2014 Nov;54(5):822–9. doi: 10.1093/icb/icu078. Epub 2014 Jun 19. PMID: 24948139.

76. Wang L, Chen P, Xiao W. β-hydroxybutyrate as an Anti-Aging Metabolite. Nutrients. 2021 Sep 28;13(10):3420. doi: 10.3390/nu13103420. PMID: 34684426; PMCID: PMC8540704.

77. Xie ZY, Zhang D, Chung DJ, Tang ZY, Huang H, et al. 2016. Metabolic regulation of gene expression by histone lysine β-hydroxybutyrylation. Mol. Cell 62:194–206

78. Yi C, Ma M, Ran L, Zheng J, Tong J, Zhu J, Ma C, Sun Y, Zhang S, Feng W, Zhu L, Le Y, Gong X, Yan X, Hong B, Jiang FJ, Xie Z, Miao D, Deng H, Yu L. 2012. Function and molecular mechanism of acetylation in autophagy regulation. Science 336(6080): 474–477. doi: 10.1126/science.1216990. PMID: 22539722.

79. Zhu X, Zhan Y, Jia X, Li M, Yin T, Wang J. Combined effects of spinetoram and Microcystis aeruginosa on Daphnia pulex offspring: Maternal effects and multigenerational implications. Chemosphere. 2024 Mar;352:141376. doi: 10.1016/j.chemosphere.2024.141376. Epub 2024 Feb 3. PMID: 38316281.

