## Supplementary Tables and Figures for "Ketone body b-Hydroxybutyrate does not extend lifespan, but upregulates fecundity in food-limited *Daphnia*, with a transgenerational effect"

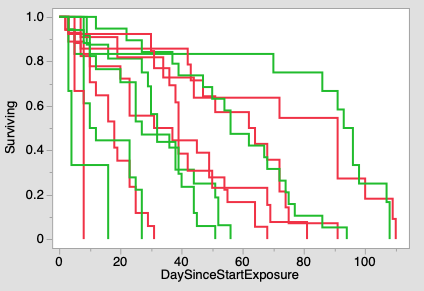

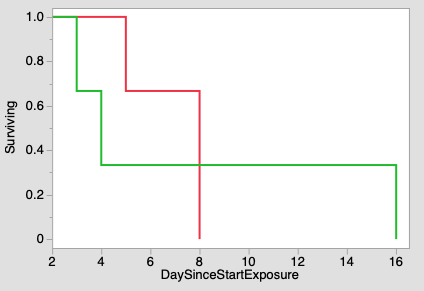

Age at start of BHB treatment: 144

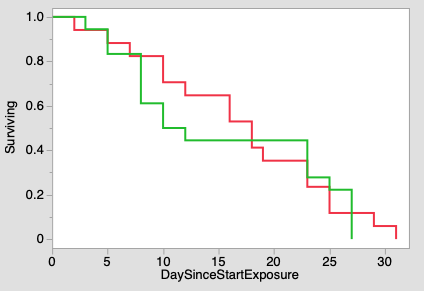

Age at start of BHB treatment: 116

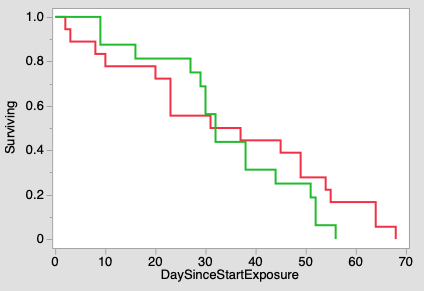

Age at start of BHB treatment: 83

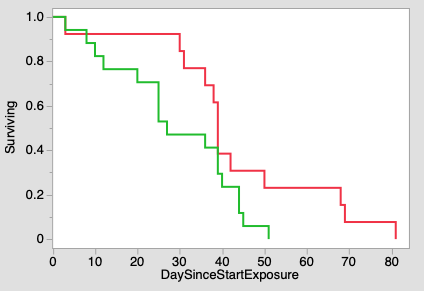

Age at start of BHB treatment: 71

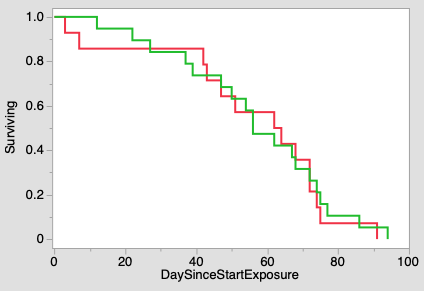

Age at start of BHB treatment: 40

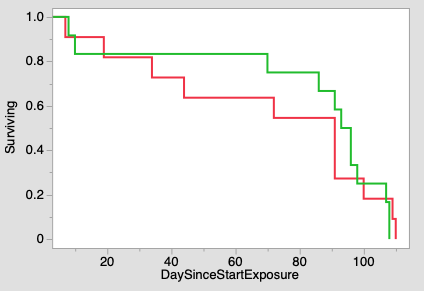

Age at start of BHB treatment: 15

Supplementary Fig.S1. Survival curves of cohorts of *Daphnia* expose to BHB for 20 days starting at different ages. Top: all cohorts shown together; right: each cohort separately. Green: control; red: BHB exposure.

Supplementary Tables

Table S1. Effects of age class (days after onset of BHB treatment), BHB exposure and clones on fecundity in Experiment 3 (REML ANOVA). Age class = Post Exposure time, in 20-day bins.

| Source | DF | | | | Sum of Squares | | | F Ratio | | | | Prob > F | | |
| --- | --- | --- | --- | --- | --- | --- | --- | --- | --- | --- | --- | --- | --- | --- |
| Age of BHB treatment onset: 83 days | | | | | | | | | | | | | | |
| BHB Treatment | | 1 | | | 4.813 | | | 7.635 | | | | 0.0062 | | |
| Age class | | 2 | | | 9.843 | | | 7.807 | | | | 0.0005 | | |
| BHB Treatment * Age class | | 2 | | | 1.038 | | | 0.823 | | | | 0.44 | | |
| Age of BHB treatment onset: 70 days | | | | | | | | | | | | | | |
| BHB Treatment | | | 1 | | | | 2.696 | | | | 2.123 | | | 0.15 |
| Age class | | | 2 | | | | 4.016 | | | | 1.581 | | | 0.21 |
| BHB Treatment * Age class | | | 2 | | | | 0.449 | | | | 0.177 | | | 0.84 |
| Age of BHB treatment onset: 40 days | | | | | | | | | | | | | | |
| BHB Treatment | | | | 1 | | 4.012 | | | 6.300 | | | | | 0.0125 |
| Age class | | | | 4 | | 34.068 | | | 13.372 | | | | | <0.0001 |
| BHB Treatment * Age class | | | | 4 | | 8.924 | | | 3.503 | | | | | 0.0080 |
| Age of BHB treatment onset: 15 days | | | | | | | | | | | | | | |
| BHB Treatment | | | 1 | | | 0.439 | | | | 1.039 | | | 0.31 | |
| Age class | | | 5 | | | 67.475 | | | | 31.916 | | | <0.0001 | |
| BHB Treatment * Age class | | | 5 | | | 9.877 | | | | 4.672 | | | 0.0003 | |

Table S2. ANCOVA of the effects of Clones, maternal BHB treatment, and neonate size at birth on lipids provisioning to neonates (median NR fluorescence). See Main text Table 3 for separate analysis of length at birth and NR fluorescence.

| Source | DF | DFDen | F Ratio | Prob > F |
| --- | --- | --- | --- | --- |
| Treatment | 1 | 43.0 | 1.317 | 0.26 |
| Clone | 2 | 46.1 | 10.098 | 0.0002 |
| Treatment*Clone | 2 | 41.8 | 4.199 | 0.022 |
| Length | 1 | 138.6 | 0.0001 | 0.99 |
